# Direct mass spectrometry-based detection and antibody sequencing of Monoclonal Gammopathy of Undetermined Significance from patient serum – a case study

**DOI:** 10.1101/2023.05.22.541697

**Authors:** Weiwei Peng, Maurits A. den Boer, Sem Tamara, Nadia J. Mokiem, Sjors P.A. van der Lans, Douwe Schulte, Pieter-Jan Haas, Monique C. Minnema, Suzan H.M. Rooijakkers, Arjan D. van Zuilen, Albert J.R. Heck, Joost Snijder

## Abstract

Monoclonal gammopathy of undetermined significance (MGUS) is a plasma cell disorder, characterized by the presence of a predominant monoclonal antibody (*i*.*e*., M-protein) in serum, without clinical symptoms. Here we present a case study in which we detect MGUS by liquid-chromatography coupled with mass spectrometry (LC-MS) profiling of IgG1 in human serum. We detected a Fab-glycosylated M-protein and determined the full heavy and light chain sequences by bottom-up proteomics techniques using multiple proteases, further validated by top-down LC-MS. Moreover, the composition and location of the Fab-glycan could be determined in CDR1 of the heavy chain. The outlined approach adds to an expanding mass spectrometry-based toolkit to characterize monoclonal gammopathies such as MGUS and multiple myeloma, with fine molecular detail. The ability to detect monoclonal gammopathies and determine M-protein sequences straight from blood samples by mass spectrometry provides new opportunities to understand the molecular mechanisms of such diseases.

## Introduction

Monoclonal gammopathy of undetermined significance (MGUS) is a plasma cell disorder, characterized by the presence of a predominant monoclonal antibody (*i*.*e*., M-protein) in patient serum. ^1^ MGUS is a preclinical stage of Multiple Myeloma (MM), with an estimated annual risk of 1% to progress to MM. ^2^ The most common antibody isotype in MGUS patients is IgG, which is a heterodimer consisting of two identical pairs of heavy chains (HC) and light chains (LC).^3,4^ All IgG share one conserved N-linked glycosylation site on each copy of the HC in the Fc region.^5^ However, the M proteins present in both MGUS and MM patients have been reported to have a high frequency of unusual additional glycosylation in the Fab region, present in the variable domains of either the light or heavy chains. ^6,7^

We recently developed methods for direct mass spectrometry-based repertoire profiling and sequencing of IgG1 from human serum.^8^ In this method IgG is affinity purified from serum samples, followed by selective digestion and release of the IgG1 Fab portion.^9^ Subsequent analysis of the released Fabs by reversed phase liquid chromatography, coupled with mass spectrometry (LC-MS), establishes a highly resolved map of antibody clones separated by mass and retention time, spanning at least 2 orders of magnitude in abundance. By spiking in monoclonal antibodies at known concentrations, also absolute concentrations of endogenous clones can be estimated by normalizing their signal intensities. Typically, we detect a few hundred of the most abundant clones, together making up 50-90% of the total subclass concentration. The most abundant IgG1 clones are generally in the order of 0.01-0.1 mg/mL, with sometimes outliers of up to about 1 mg/mL in hospitalized patients in critical condition.^10,11^

During screening of serum IgG1 repertoires of a new cohort of donors by LC-MS we observed a donor whose repertoire was exceptionally dominated by a single IgG1 clone exhibiting a very high concentration of approximately 10 mg/mL in serum. This single clone contributes approximately 98% to the total amount of all IgG1 molecules in the serum of this patient. Subsequent clinical tests confirmed diagnosis of MGUS. Our MS data also indicated that this MGUS M-protein harbored abundant Fab glycosylation. Combining the above IgG1 profiling method with bottom-up proteomics-based sequencing, we were able to recover the full sequence of the antibody heavy and light chains. The Fab glycosylation could be traced back to a specific residue in CDR1 of the heavy chain. The attached N-glycan structures could be assigned and quantified based on the intact Fab MS spectra and tandem MS spectra of the corresponding glycopeptides. This case illustrates how integrated bottom-up and top-down proteomics can be used to detect MGUS and other monoclonal gammopathies, sequence the associated monoclonal antibody against a background of serum IgG1, even when the M-protein is Fab-glycosylated, and that the composition of this Fab-glycosylation can be determined and localized from the same sampled material.

## Results

### Observation of a Fab-glycosylated IgG1 M protein

When analyzing the serum IgG1 clonal repertoire of a patient that had undergone a recent kidney transplant, we unexpectedly encountered an atypical antibody profile. This patient was part of a longitudinal study cohort who were observed after kidney transplantation for immunological monitoring. Per protocol in these patients, serum samples were obtained at different time points, starting from moments before surgery (t = 0 days) with follow-up samplings for over a year which were stored in a biobank. Samples of patients who developed designated bacterial infections in follow up were drawn from the biobank and to investigate how the antibody repertoire responded to the surgery and subsequent infections, we applied our LC-MS IgG1 profiling approach to these longitudinal serum samples.

Strikingly, the IgG1 repertoire of this particular kidney transplant patient was very different from that of other donors, being dominated by seemingly a few extremely abundant clones that also exhibit relatively high masses (Figure 1A). This pattern remained unchanged before or after kidney transplantation or even after an episode of sepsis with a *Klebsiella* species (Supplementary Figure S1).

**Figure 1.**
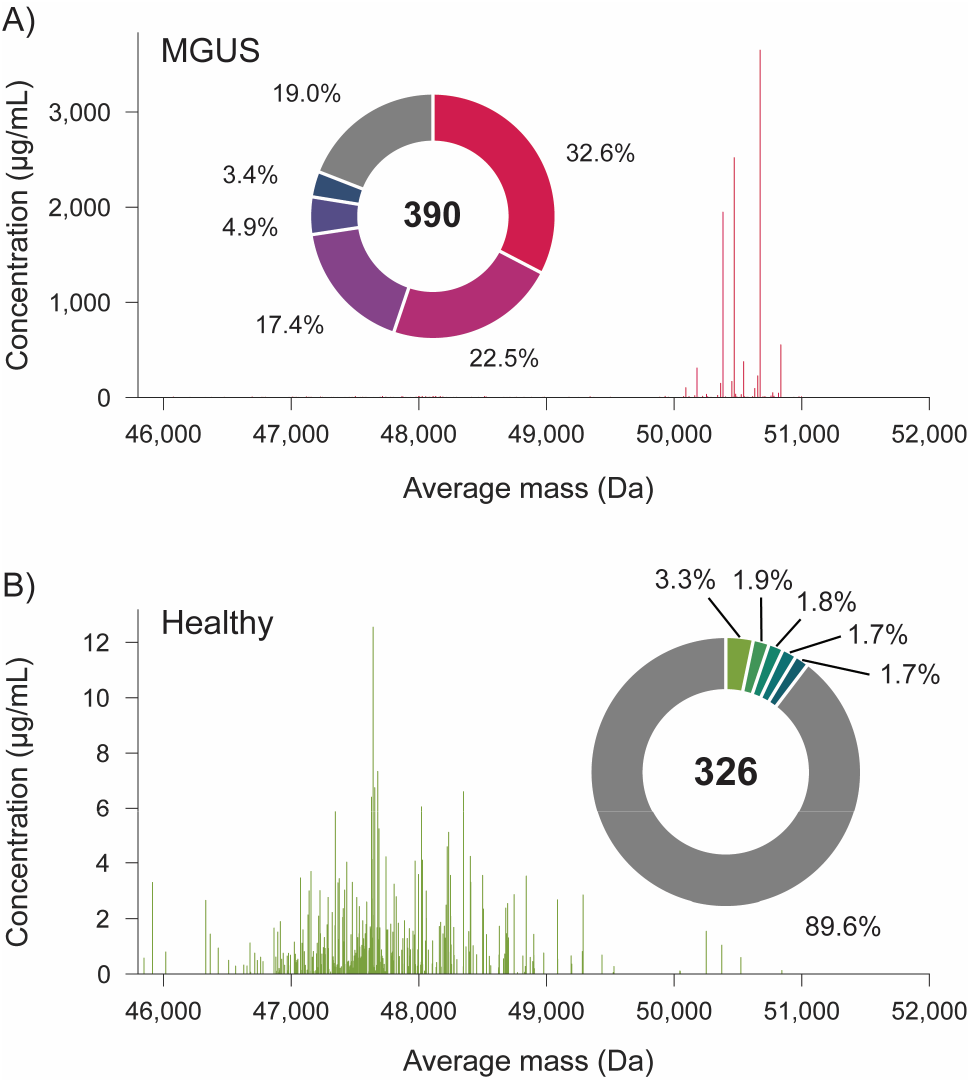
Detection of MGUS by LC-MS based IgG1 Fab-profiling. A) Serum IgG1 Fab profile of human subject with putative MGUS. B) Illustrative IgG1 Fab profile from serum of a healthy human donor. Note the difference in complexity, average mass, and concentration of detected IgG1 Fabs, hinting at the presence of a Fab-glycosylated M-protein in A). Pie charts show the relative abundance (%) of the top-5 most abundant Fab species.

**Figure 2.**
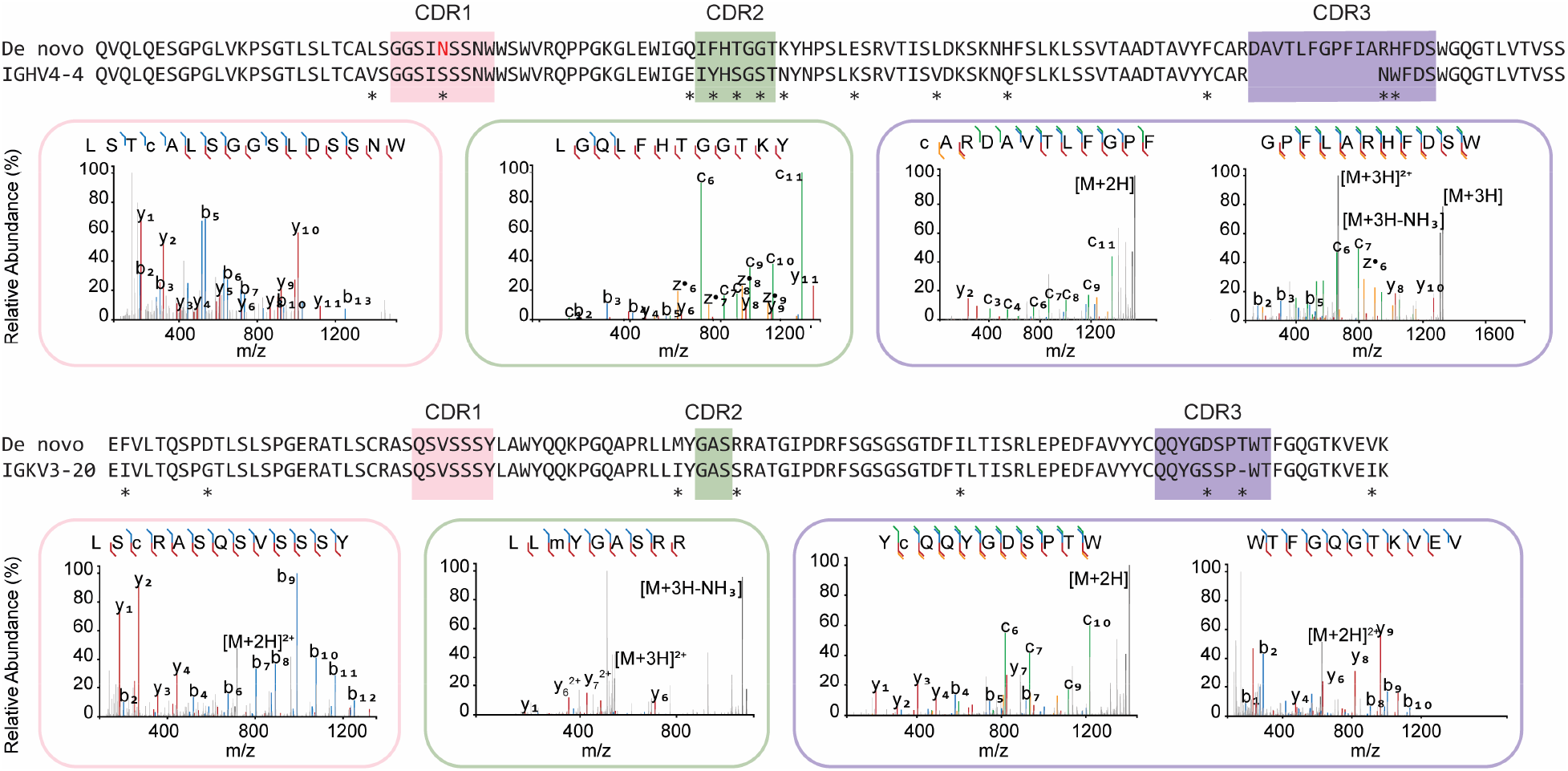
De novo sequencing of the MGUS Fab by bottom-up proteomics. The variable region alignment to the inferred germline sequence is shown for both heavy and light chains. Positions with putative somatic hypermutation are highlighted with asterisks (*). The MS/MS spectra supporting the annotation of the CDRs are shown beneath the sequence alignment. b/y ions are indicated in blue and red, while c/z ions are indicated in green and yellow.

Because these antibodies were detected at relatively high masses, we hypothesized that the unusual antibody profile of the patient resulted from a single IgG1 clone of very high concentration that may carry Fab-glycosylation. Based on experimental data from our lab and theoretical calculations using sequences from the ImMunoGeneTics (IMGT) database,^12^ the bulk of IgG1 Fabs have backbone masses of roughly 46-50 kDa. The abundant Fabs that we detected, however, have higher masses of more than 50 kDa. This strongly suggests that they are modified by N-glycosylation, which would contribute roughly 2 kDa to their mass. Combining the signal intensities of the multiple putative glycoforms, the total concentration of this clone is approximately 10 mg/mL at t = 0, remaining high throughout the longitudinal follow up (see Supplementary Figure S1). Compared to the total IgG1 concentration in human plasma, approximately 8 mg/mL on average, this is extremely high, prompting additional clinical tests. The patient was tested for an M protein using serum immunofixation, which confirmed the IgG kappa M protein, as well as serum electrophoresis, which could not quantify the M protein due to low amount. In addition, free light chains (FLC) were determined (Binding Site®), demonstrating an elevated FLC kappa of 61.52 mg/L, and a FLC ratio of 3.92 (normal range 0.26-1.65). No bone marrow biopsy was done and a diagnosis of monoclonal gammopathy of undetermined significance (MGUS) was made.

### Direct MS-based sequencing and glycan localization of the serum-derived MGUS clone

We have recently demonstrated the direct MS-based sequencing of serum-derived antibodies, using bottom-up proteomics methods.^2^ Owing to the high abundance of the MGUS M-protein in the serum samples of this donor, we were able to sequence the full antibody without further fractionation. First, we used an in-gel digestion protocol, using in parallel four proteases of complementary specificity, to obtain overlapping peptides for *de novo* sequencing by LC-MS/MS analysis. We identified the M-protein as consisting of an IGHV4-4 heavy chain, coupled with an IGKV3-20 light chain. Notably, this experiment was performed with intact N-glycosylation and resulted in a lack of coverage in CDR1 of the heavy chain. The germline sequence of IGHV4-4 CDRH1 contains 5 serine residues and a single asparagine, priming it to obtain an N-glycosylation sequon by as many as 6 independent substitutions. These observations pointed to CDRH1 as a likely region to contain the putative Fab-glycosylation.

Digestion with PNGase F results in the removal of the N-glycan and converts the glycan-linked asparagine to an aspartic acid residue.^13^ We digested the serum sample with PNGase F, followed by proteolysis with chymotrypsin and thermolysin in parallel. This recovered the previously missing CDRH1 sequence, containing a clear DSS motif, which would have corresponded with an NSS glycosylation sequon in the antibody prior to PNGase F digestion. Using the experimentally determined sequence, we then performed a glycoproteomics database search including common human N-glycans and were able to detect a predominant HexNAc(5)Hex(5)Fuc(1)NeuAc(2) glycan at the identified NSS sequon in CDRH1 (see Supplementary Figure S2).

### Validation of the sequence and Fab glycosylation by middle-down LC-MS/MS

To validate our bottom-up *de novo* sequencing result, we further analyzed the MGUS Fab by native MS and middle-down LC-MS/MS. The intact mass profile of the Fab is shown in Figure 3A. The observed masses are consistent with the determined sequence, considering a pyroglutamic acid modification of the heavy chain N-terminus, and reveals heterogeneous glycosylation with a predominant HexNAc(5)Hex(5)Fuc(1)NeuAc(2) glycan, as also observed by bottom-up LC-MS/MS (see Supplementary Table S1). A predicted structural model of the variable domain shows this glycan protruding outwards from CDRH1, leaving the other CDR loops exposed (see Figure 3B). The observed Fab glycosylation pattern follows a similar trend as reported by Bondt *et al*. in that it is enriched in galactosylation, sialylation, and bisection compared to Fc glycosylation at the conserved N297 site (see Figure 3C; a full overview of glycoforms of the MGUS Fab is provided in Supplementary Table S1).^14^

**Figure 3.**
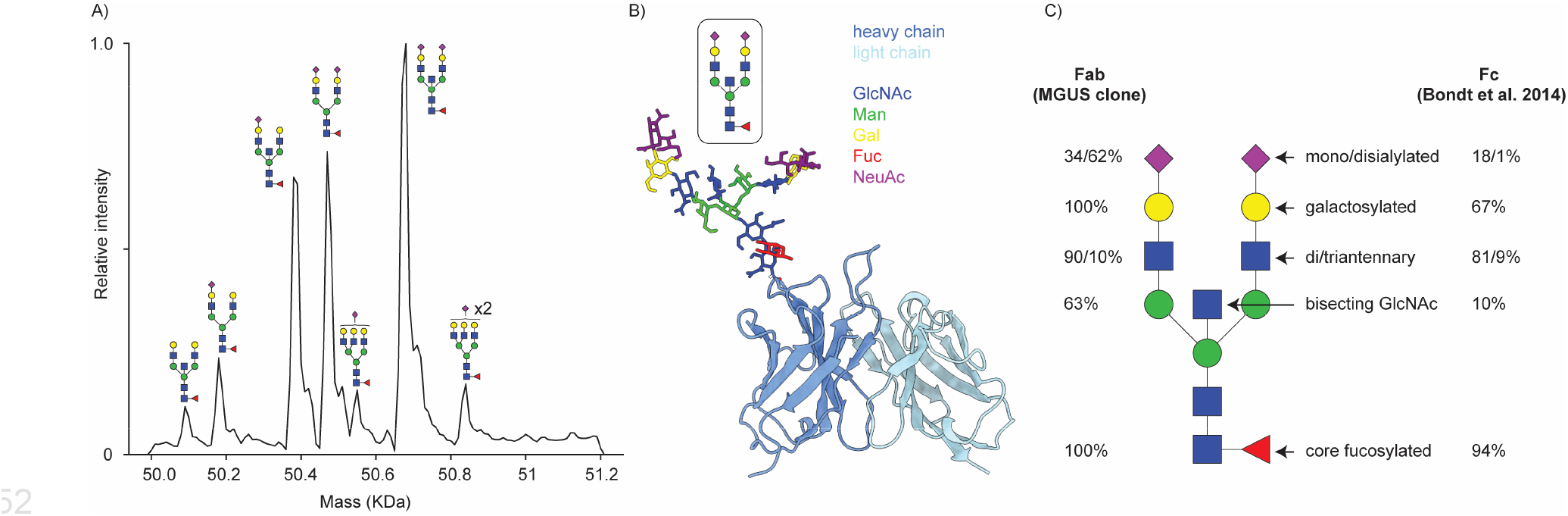
N-Glycosylation of MGUS Fab. A) Intact mass profile from native MS; peaks are annotated according to the assigned glycan structure. B) ABodyBuilder2 structural model prediction of the MGUS Fab variable domain with HexNAc(5)Hex(5)Fuc(1)NeuAc(2) grafted on CDRH1 using GLYCAM. C) Glycosylation profile of MGUS Fab compared to typical Fc glycosylation at N297, according to Bondt et al. 2014.

We performed middle-down fragmentation of the reduced Fab by EThcD to confirm the sequence determined by bottom-up proteomics. This resulted in a coverage of 25.7% for the Fd and 60.5% for the LC (see Figure 4 and Supplementary Figure S3). We attribute the lower coverage of the Fd to the presence of the N-glycan in CDRH1, which is likely to fragment during EThcD, producing more complex spectra, with lower signal for individual fragments. Furthermore, glycan fragmentation is not yet implemented in currently available peak matching algorithms for middle-down LC-MS/MS. Nonetheless, the intact masses and middle-down fragmentation patterns support the M-protein sequence determined by bottom-up proteomics.

**Figure 4.**
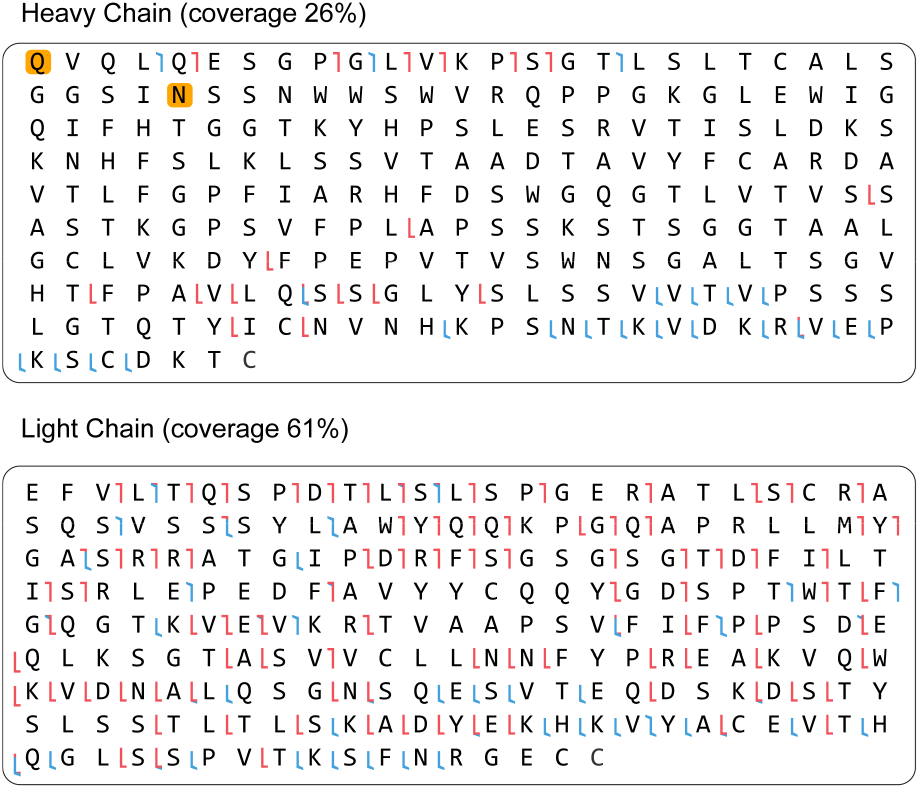
Top-down LC-MS/MS data on the of Fab-glycosylated M-protein. Shown are the heavy and light chain sequences, position supported by b/c or y/z ions from EThcD fragmentation are indicated in red and blue, respectively.

## Conclusion

Here we demonstrate that LC-MS based IgG1 profiling of patient serum can lead to the detection of an M-protein, which can be related to diseases such as MGUS or Multiple Myeloma. The associated sequence of the M-protein can be fully derived by mass spectrometry. In this case, mass spectrometry also revealed the presence, location and composition of Fab glycosylation in the heavy chain of the M-protein. The outlined approach adds to an expanding mass spectrometry-based toolkit to characterize monoclonal gammopathies such as MGUS and MM with fine molecular detail.^15–20^ The ability to detect monoclonal gammopathies and determine M-protein sequences straight from peripheral blood samples by mass spectrometry provides opportunities to understand the molecular mechanisms of these diseases.

## Materials and Methods

### Cohort and Trial information

In the period of 2015-2019 patients who underwent kidney transplantation were asked to participate in a biobank to evaluate immunological developments after kidney transplantation. All participants provided written informed consent to collect clinical data and serum samples pre-transplantation and at month 1, 3, 6, and 12 post-transplantation. The study was approved by the local Biobank Research Ethics Committee (protocol 15-019). Serum samples of patients with a recorded bacterial infection after kidney transplantation were identified and analyzed to evaluate the immunological response to such an infection. In one of the patients the samples showed a few extremely abundant clones.

### IgG purification and Fab production

IgG1 clonal profiling was performed based on a method previously described by Bondt et al. ^8^ Two internal reference mAbs (trastuzumab and alemtuzumab) were spiked into serum samples of 10 μL to a final concentration of 20 μg/mL (200 ng), after which IgG was captured using 10 μL CaptureSelect FcXL affinity matrix (20 μL slurry, Thermo Fisher) in a spin column. After binding for 60 min on a shaker at 750 rpm and room temperature, columns were washed in four sequential rounds by adding 150 μL PBS and removing the liquid by centrifugation for 1 min at 500 g. IgG1 Fab molecules were released on through on-bead proteolytic digestion using 100 U IgdE (FabALACTICA; Genovis) in 50 μL 150 mM sodium phosphate pH 7.0 overnight on a shaker at 750 rpm and 37 °C. Liquid containing free IgG1 Fabs was captured through centrifugation for 1 min at 1,000 g.

Analysis was performed by reversed-phase LC-MS using a Vanquish Flex UHPLC system (Thermo Fisher) coupled to an Orbitrap Exploris 480 instrument (Thermo Scientific. Chromatographic separation was performed on a 1 × 150 mm MAbPac column at 80 °C and using a flow rate of 150 μL/min. Mobile phase A consisted of MilliQ water with 0.1% formic acid, mobile phase B of acetonitrile with 0.1% formic acid. Samples were run starting with a 10%-25% B ramp with the spray voltage turned off for 2 minutes to wash away salts. This was followed by a 54 min linear gradient of 25%-40% B, a 95% B wash and re-equilibration at 10% B. The mass spectrometer was operated at low pressure setting in Intact Protein mode at a set resolution of 7,500 at 200 *m/z*. For every scan, 5 μscans were acquired with an *m/z* range of 500 - 4,000 using an AGC target of 300% with a maximum injection time of 50 ms. The RF lens was set to 40% and a source fragmentation energy of 15 V was used. Raw data were processed by sliding window deconvolution using the ReSpect algorithm in BioPharma Finder v3.2 (Thermo Fisher). Further analysis was performed using an in-house python library described by Bondt et al. Components with masses between 45,000 and 53,000 Da, most intense charge states above m/z 1,000, and a Score of over 40 were considered valid Fab identifications.

### Bottom-up proteomics

#### In-gel digestion

Fab (3 μg/lane) was loaded on a 4%-12% Bis-Tris precast gel (Bio-rad) in non-reducing conditions and run at 120 V in 3-Morpholinopropane-1-sulfonic acid (MOPS) buffer (Bio-rad). Bands were visualized with Imperial Protein Stain (Thermo Fisher Scientific), and the size of the fragments evaluated by running a protein standard ladder (Bio-rad). The Fab bands were cut and reduced by 10 mM TCEP at 37°C, then alkylated in 40 mM IAA at RT in the dark, followed by alkylation in 40 mM IAA at RT in the dark. The Fab bands were digested by trypsin, chymotrypsin, thermolysin, and alpha lytic protease at 37 °C overnight in 50 mM ammonium bicarbonate buffer. The peptides were extracted with two steps incubation at RT in 50% ACN, and 0.01% TFA, and then 100% ACN respectively. The peptides were dried in speed-vac. To obtain the sequence of the glycosylated Fab, the N-linked glycan was removed by PNGaseF at 37 °C overnight then in gel digested as described above.

#### Mass Spectrometry

The digested peptides were separated by online reversed phase chromatography on an Dionex UltiMate 3000 (Thermo Fisher Scientific) (column packed with Poroshell 120 EC C18; dimensions 50 cm × 75 μm, 2.7 μm, Agilent Technologies) coupled to a Thermo Scientific Orbitrap Fusion mass spectrometer or Thermo Scientific Orbitrap Fusion LUMOS mass spectrometer. Samples were eluted over a 90 min gradient from 0 to 35% acetonitrile at a flow rate of 0.3 μL/min. Peptides were analyzed with a resolution setting of 60 000 in MS1. MS1 scans were obtained with a standard automatic gain control (AGC) target, a maximum injection time of 50 ms, and a scan range of 350–2000. The precursors were selected with a 3 m/z window and fragmented by stepped high-energy collision dissociation (HCD) and electron-transfer higher-energy collision dissociation (EThcD). The stepped HCD fragmentation included steps of 25, 35, and 50% normalized collision energies (NCE). EThcD fragmentation was performed with calibrated charge-dependent electron-transfer dissociation (ETD) parameters and 27% NCE supplemental activation. For both fragmentation types, MS2 scans were acquired at a 30 000 resolution, a 4e5 AGC target, a 250 ms maximum injection time, and a scan range of 120–3500.

#### Peptide Sequencing from MS/MS Spectra

MS/MS spectra were used to determine *de novo* peptide sequences using PEAKS Studio X (version 10.6).^21,22^ We used a tolerance of 20 ppm and 0.02 Da for MS1 and 0.02 Da for MS2, respectively. Carboxymethylation was set as fixed modification of cysteine and variable modification of peptide N-termini and lysine. Oxidation of methionine and tryptophan, pyroglutamic acid modification of N-terminal glutamic acid, and glutamine were set as additional variable modifications. The CSV file containing all the *de novo* sequenced peptides was exported for further analysis.

#### Template-based assembly via Stitch

Stitch^23^ (1.1.2) was used for the template-based assembly. The human antibody database from IMGT was used as template. The cutoff score for the *de novo* sequenced peptide was set as 90/70 and the cutoff score for the template matching was set as 10. All the peptides supporting the sequences were examined manually. The ions for annotation of the CDR regions were exported and visualized by Interactive Peptide Spectral Annotator.^24^

#### Glycoproteomics data analysis

Chymotryptic digested peptides were used to search for site specific glycosylation via Byonic (v5.0.3).^25^ The *de novo* obtained sequences were selected as protein database. Four missed cleavages were permitted using C-terminal cleavage at WFMLY for chymotrypsin. Carboxymethylation of cysteine was set as fixed modification, oxidation of methionine/tryptophan as variable rare 1, Gln to pyro-Glu and Glu to pyro-glu on the N-temmius of protein as rare 1, and N-glycan modifications were set as variable rare 1. The N-glycan 132 human database from Byonic was applied in the search. All reported glycopeptides in the Byonic result files were manually inspected for quality of fragment assignments.

### Native MS

To remove the N-linked glycan on the fab, the samples was incubated with 1% Rapigest (Waters Corporation, USA) for 3 min at 90 °C.the PNGaseF was added to the sample and incubated at 50 °C for 10 min. Both the native fab and the deglycosylated fab were buffer exchanged into 150 mM ammonium acetate (pH 7.5) using Amicon 10 kDa MWCO centrifugal filters (Merck Millipore). The samples were loaded into gold-coated borosilicate capillaries (in-house prepared) and analyzed on an ultra-high mass range (UHMR) Q-Exactive Orbitrap (Thermo Fisher Scientific, Bremen, Germany). The mass spectra were obtained in positive mode with an ESI voltage of 1.3 kV. The maximum injection was set at 100 ms and the HCDenergy was set at 100 V. The used resolution was 12500 at 400 m/z. The S-Lens level was set at 200. UniDec was used to generating the charge-deconvoluted spectrum. ^26^

### MD proteomics

The reduced Fab was freshly prepared by incubating with TCEP at 60°C for 30 min before injecting to MS. Around 1 μg sample was used for a single measurement. Reduced Fab was measured by LC-MS/MS. Samples were loaded on a Thermo Scientific Vanquish Flex UHPLC instrument, equipped with a 1 mm x 150 mm MAbPac RP analytical column, directly coupled to an Orbitrap Fusion Lumos Tribrid (Thermo Scientific, San Jose, CA, USA). The samples were eluted over 22 min at a 150 μL/min flow rate. Gradient elution was achieved by using two mobile phases A (0.1% HCOOH in Milli-Q) and B (0.1% HCOOH in CH3CN) and ramping up B from 10 to 25% over one minute, from 25 to 40% over 14 min, and from 40 to 95% over one minute. MS data were collected with the instrument operating in Intact Protein and Low Pressure mode. Spray voltage was set at 3.5 kV, capillary temperature 350 °C, probe heater temperature 100 °C, sheath gas flow 15, auxiliary gas flow 5, and source-induced dissociation was set at 15 V. Separate Fab chains were analyzed with a resolution setting of 120,000. MS1 scans were acquired in a range of 500-3,000 Th with the 250% AGC target and a maximum injection time set to either 50 ms for the 7,500 resolution or 250 ms for the 120,000 resolution. In MS1, 2 μscans were recorded for the 7,500 resolution and 5 μscans for the 120,000 resolution per scan. Data-dependent mode was defined by the number of scans: single scan for intact Fabs and two scans for separate Fab chains. MS/MS scans were acquired with a resolution of 120,000, a maximum injection time of 500 ms, a 1,000% AGC target, and 5 μscans averaged and recorded per scan for the separate Fab chains. The EThcD active was set at true. The ions of interest were mass-selected by quadrupole in a 4 Th isolation window and accumulated to the AGC target prior to fragmentation. MS/MS spectra were used to validate the sequences using LC-MS Spectator (Version 1.1.8313.28552) and ProSight Lite (1.4.8).^27,28^ In LC-MS Spectator, we used a tolerance of 10 ppm for MS1 and 20 ppm for MS2, respectively and applied the S/N threshold filtering (1.5). All the annotated ions were exported and visualized in ProSight Lite.

### Structural model of glycosylated MGUS Fab

The variable domain of the MGUS Fab was modelled using the ABodyBuilder2 webserver from the SAbPred suite. The predominant HexNAc(5)Hex(5)Fuc(1)NeuAc(2) glycoform was modelled as diantennary, bisected complex glycan with core fucosylation using the GLYCAM glycoprotein builder webserver. Figures were rendered in ChimeraX.

## Acknowledgements

This research was funded by the Dutch Research Council NWO Gravitation 2013 BOO, Institute for Chemical Immunology (ICI; 024.002.009) to J.S and through the NWO TTW-NACTAR Grant #16442 (to A.J.R.H. and S.H.M.R.). We additionally acknowledge support from NWO, funding the MS facilities through the X-omics Road Map program (project 184.034.019).

## Data availability

The raw LC-MS/MS files and analyses have been deposited to the ProteomeXchange Consortium via the PRIDE partner repository with the dataset identifier PXD042266.

