## Supplementary Information for "Direct mass spectrometry-based detection and antibody sequencing of Monoclonal Gammopathy of Undetermined Significance from patient serum – a case study"

*Supplementary Table S1. Intact mass analysis of MGUS Fab and derivatives. Glycoforms are denoted with: N-acetyl hexosamine (N), hexose (H), fucose (F), and N-acetyl neuraminic acid (S).*

| **Sample** | **glycoform** | **mass exp. (Da)** | **SD (Da)[c]** | **mass theor. (Da)[d]** | **Δmass (Da)** | **relative intensity (%)** |
|---|---|---|---|---|---|---|
| intact Fab[a] | N5H5F1 | 50093.96 | 0.40 | 50090.79 | 3.17 | 3.63 |
| | N4H5F1S1 | 50181.62 | 0.16 | 50178.85 | 2.77 | 7.52 |
| | N5H5F1S1 | 50384.96 | 0.21 | 50382.05 | 2.91 | 22.04 |
| | N4H5F1S2 | 50472.65 | 0.02 | 50470.11 | 2.54 | 23.78 |
| | N5H6F1S1 | 50549.69 | 0.04 | 50544.19 | 5.50 | 4.93 |
| | N5H5F1S2 | 50675.96 | 0.13 | 50673.30 | 2.66 | 32.57 |
| | N5H6F1S2 | 50837.75 | 0.17 | 50835.45 | 2.30 | 5.54 |
| Fab + PNGase F[a] | - | 48120.43 | 0.12 | 48117.67 | 2.76 | - |
| Fab + TCEP HC[b] | N5H5F1S2 | 26952.69 | 0.37 | 26951.09 | 1.60 | - |
| Fab + TCEP LC[b] | - | 23729.78 | 0.37 | 23731.31 | -1.53 | - |

[a] *native MS*

[b] *LC-MS*

[c] *average and standard deviations are calculated across the series of charge states for native MS, across 4 replicate measurements for LC-MS.*

[d] *average theoretical mass considering disulfide bond formation and pyroglytamic acid conversion of the heavy chain N-terminus.*

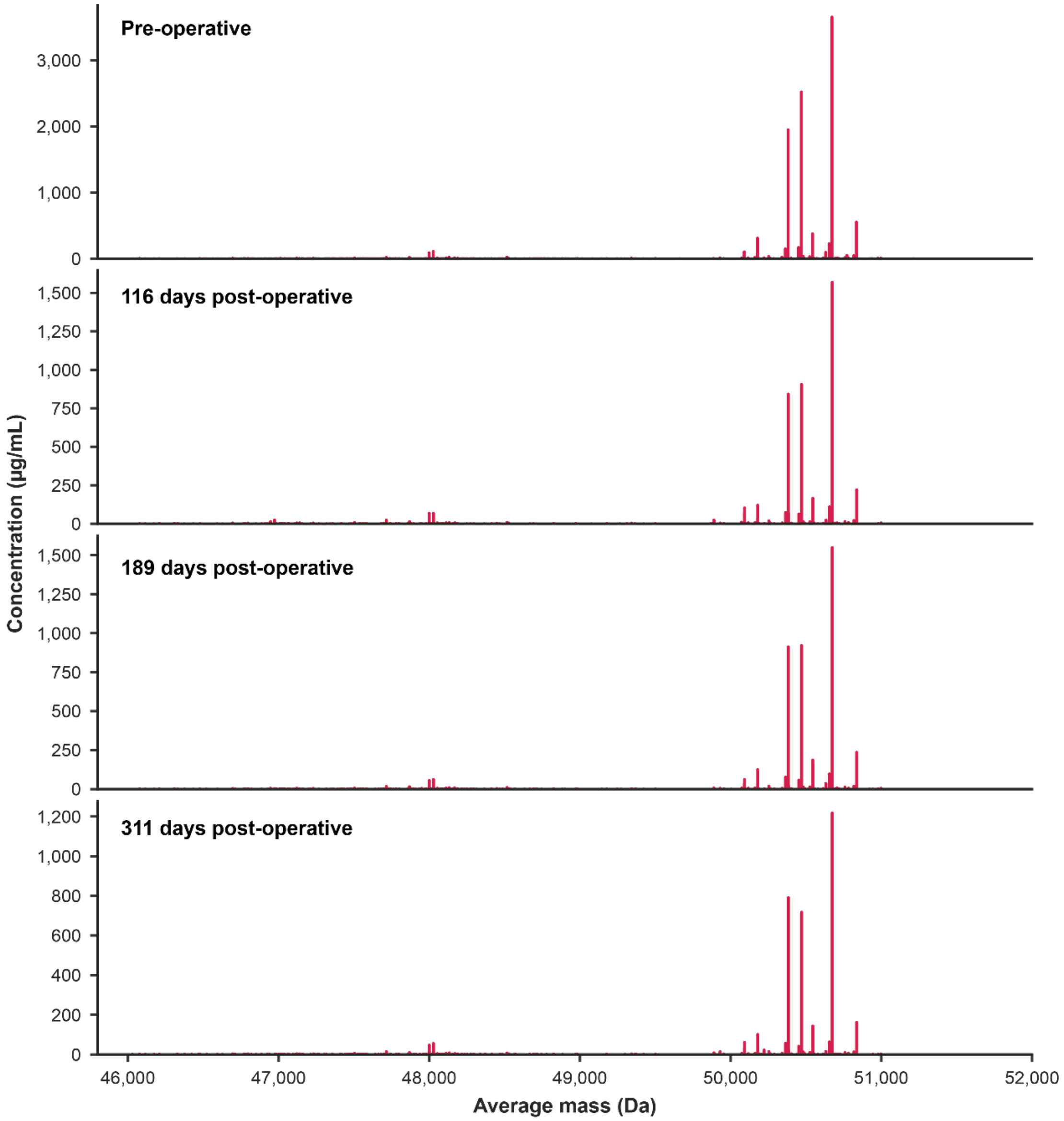


*Supplementary Figure S1. Longitudinal IgG1 Fab profile of the MGUS patient.*

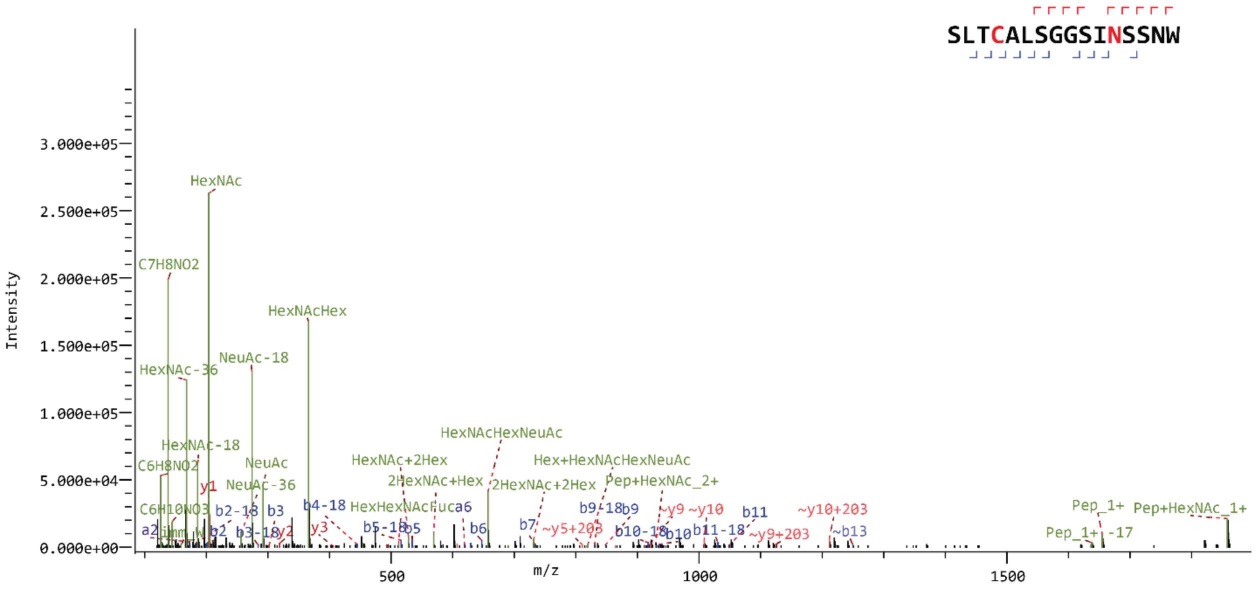


*Supplementary Figure S2. Tandem MS spectrum of N-glycopeptide in CDRH1 of M-protein. Shown is the HexNAc(5)Hex(5)Fuc(1)NeuAc(2) glycoform.*

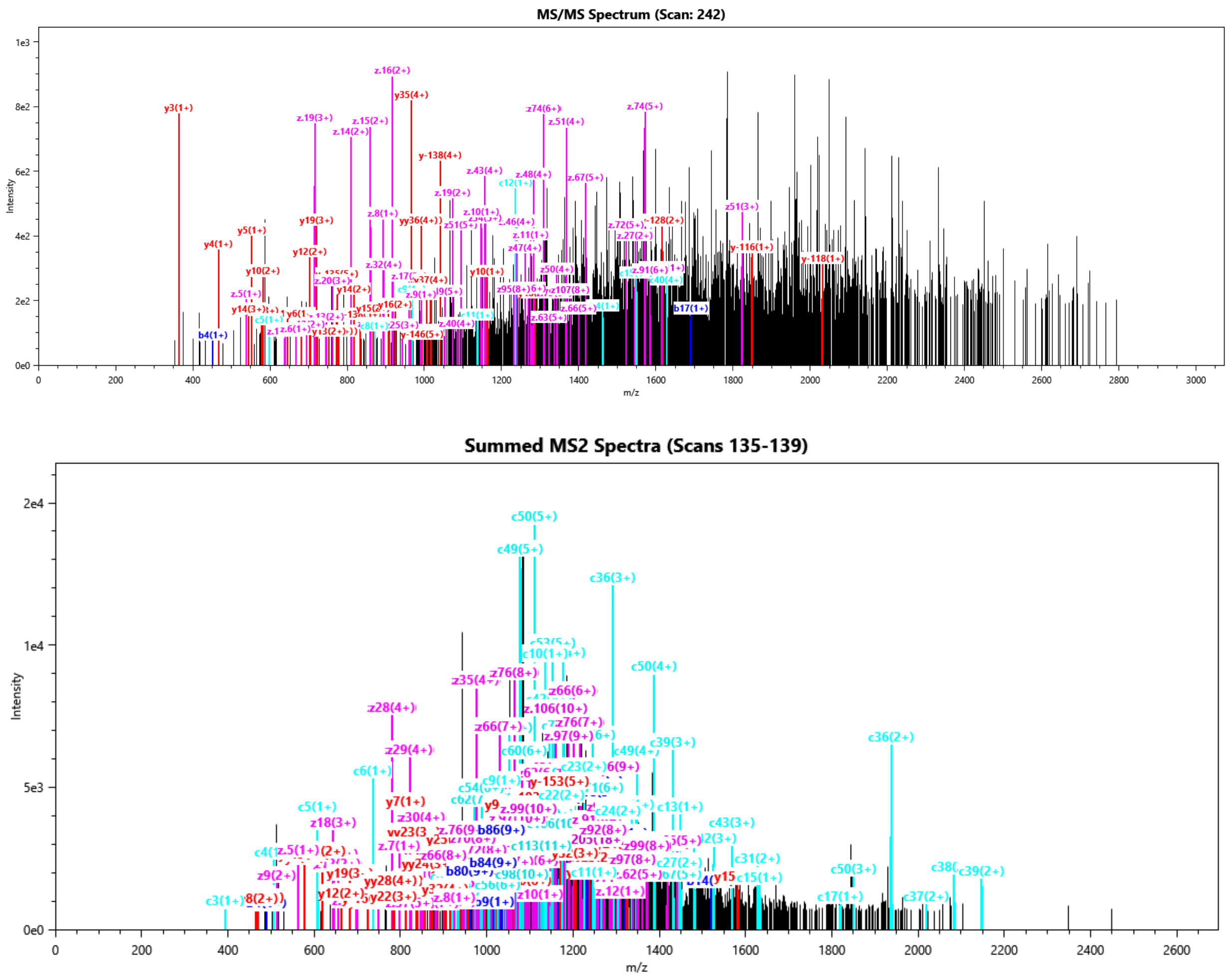


*Supplementary Figure S3. Top-down LC-MS/MS spectra of M-protein Fab Heavy Chain (top) and Light Chain.*
